# Astrocytic CREB regulates transcriptional, neuronal, and behavioral responses to cocaine

**DOI:** 10.1101/2025.06.04.657899

**Authors:** Leanne M. Holt, Angelica Minier-Toribio, Rita Futamura, Caleb J. Browne, Freddyson J. Martínez-Rivera, Tamara Markovic, Trevonn M. Gyles, Szu-ying Yeh, Eric M. Parise, Matthew Rivera, Corrine Azizian, Yun Young Yim, Veronika Kondev, Molly Estill, Shi Yan, Ellie Keane, Alexa Labanca, Giselle Rojas, Yan Dong, Eric J. Nestler

## Abstract

Drug addiction is characterized by neuronal adaptations that support a shift from goal-directed behaviors to habitual, compulsive drug-seeking with persistent effects on cognition and decision-making. Emerging evidence increasingly indicates that astrocytes are also involved in nervous system disorders, including addiction, but the cocaine-induced astrocyte-specific transcriptome has not yet been investigated. We utilized whole cell sorting of astrocytes, RNA-sequencing, and bioinformatic approaches to characterize the astrocyte transcriptome in the nucleus accumbens (NAc), a key brain region involved in reward-processing, following cocaine self-administration, prolonged abstinence, and “relapse” in male mice. We found that astrocytes exhibit robust and contextually-specific transcriptional signatures that converge strongly with human cocaine use disorder. Bioinformatic analysis revealed CREB as a highly ranked predicted upstream regulator of cocaine-induced transcriptional regulation in NAc astrocytes, and CUT&RUN-sequencing mapped increased CREB binding across the astrocyte genome in response to cocaine. Viral-mediated manipulation of CREB activity selectively in NAc astrocytes, in combination with several measures of addiction-related behaviors including conditioned place preference and self-administration, revealed that astrocytic CREB increases the rewarding and reinforcing properties of cocaine. This effect is sex-specific, with no change in astrocytic CREB activity or CPP found in females. Subsequent experiments identify potential molecular mechanisms of astrocytic CREB’s influence through modulating astrocytic Ca^2+^ signaling in response to cocaine. Finally, we show that astrocytic CREB selectively modulates D1-type medium spiny neurons in NAc to control cocaine-related behaviors. Together, these data demonstrate that the astrocyte transcriptome responds robustly to cocaine and that CREB mediates cocaine’s effects on gene expression in astrocytes, with consequent effects on neuronal activity and rewarding responses to the drug.

## Introduction

Substance use disorders (SUDs) remain a devastating contributor to the US public health burden, with recent data demonstrating a dramatic rise in overdose deaths, including those involving cocaine use^1^. Astrocytes—the most abundant glial cell type—are increasingly implicated in disorders of the nervous system, including cocaine use disorder (CUD)^2–4^.

Within the nucleus accumbens (NAc), a key region for reward learning and motivated behaviors, astrocytes not only functionally respond to cocaine but also mediate addiction-related behaviors. Ex-vivo imaging reveals astrocytes increase calcium dynamics in response to acute cocaine, while chronic exposure decreases their calcium transients^5–7^. Enhancing astrocyte calcium signaling within the NAc through Gq-Designer Receptors Exclusively Activated by Designer Drugs (DREADDs) attenuates cocaine seeking, whereas reducing calcium dynamics via hPMCA2 overexpression in the dorsal striatum increases seeking behavior during cued reinstatement in rats^8^. Extinction training following cocaine self-administration in rodents reduces astrocyte morphological complexity and synaptic enwrapment in the NAc, while astrocyte-derived thrombospondins mediate cocaine-seeking through the formation of silent synapses^7,9,10^. Moreover, modulation of astrocytic function throughout the mesolimbic pathway further demonstrates their role in addiction-related behaviors. For example, blocking astrocytic lactate release in the basolateral amygdala during retrieval reduces cocaine seeking in rats, and astrocytic modulation of tonic inhibition in ventral tegmental area GABAergic neurons enhances cocaine’s rewarding effects ^11,12^.

Investigations of transcriptional responses to drugs of abuse reveal lasting changes in gene expression throughout the brain’s reward circuitry, thought to mediate the underlying pathophysiology of addiction ^13–17^. Despite increasing evidence for functional astrocytic contributions to addiction-related behaviors, few publications have examined the astrocyte-specific transcriptome in response to cocaine self-administration (SA), and none have investigated underlying astrocytic transcriptional mechanisms that may mediate their influence of addiction-related behaviors (SA)^18–21^. We therefore performed RNA-sequencing (RNA-Seq) on NAc astrocytes from male mice after cocaine SA, including abstinent and “relapse” conditions.

Our results revealed a robust, context-dependent transcriptional response implicating cAMP Response Element-Binding Protein (CREB) as a key cocaine-induced astrocytic transcriptional regulator. Cleavage Under Targets and Release Using Nuclease (CUT&RUN) identified putative cocaine-induced gene targets of astrocytic CREB, with a direct target involved in astrocyte calcium dynamics. We therefore probed if astrocytic CREB alters cocaine-induced calcium transients, and further demonstrate astrocytic CREB selectively increases the activity of D1-type medium spiny neurons (D1-MSNs). Viral-mediated manipulations coupled with addiction-related behavioral paradigms demonstrated that astrocytic CREB increases the rewarding and reinforcing effects of cocaine. Finally, integrating astrocytic CREB manipulations with neuronal-specific Gi-DREADDs, we established a mechanistic link between astrocytic transcriptional regulation and the neuronal responses underlying cocaine reward and reinforcement.

## Results

### Cocaine-induced regulation of the astrocyte transcriptome in the NAc

To model aspects of CUDs, male mice self-administered saline or cocaine for 10 days (d) (0.5 mg/kg infusions; 2-hour (hr) sessions; Fig 1A). Mice self-administering cocaine, but not saline, discriminated between the active and inactive levers, and received more infusions compared to saline controls (Fig 1B). Following 30 d of forced abstinence, mice were randomly assigned to receive a subcutaneous injection of saline (saline+saline [SS] and cocaine+saline [CS]) or cocaine (saline+cocaine [SC] and cocaine+cocaine [CC]) and placed back in the operant boxes for a drug-seeking test (30 minutes [min]). Mice assigned to the withdrawal condition (SW and CW) were taken directly from their homecage. This design was used previously to model aspects of addiction, including withdrawal (CW), initial drug exposure (SC), drug context re-exposure without drug (CS), and “relapse”—context re-exposure with drug (CC) ^15,22^.

**Figure 1.**
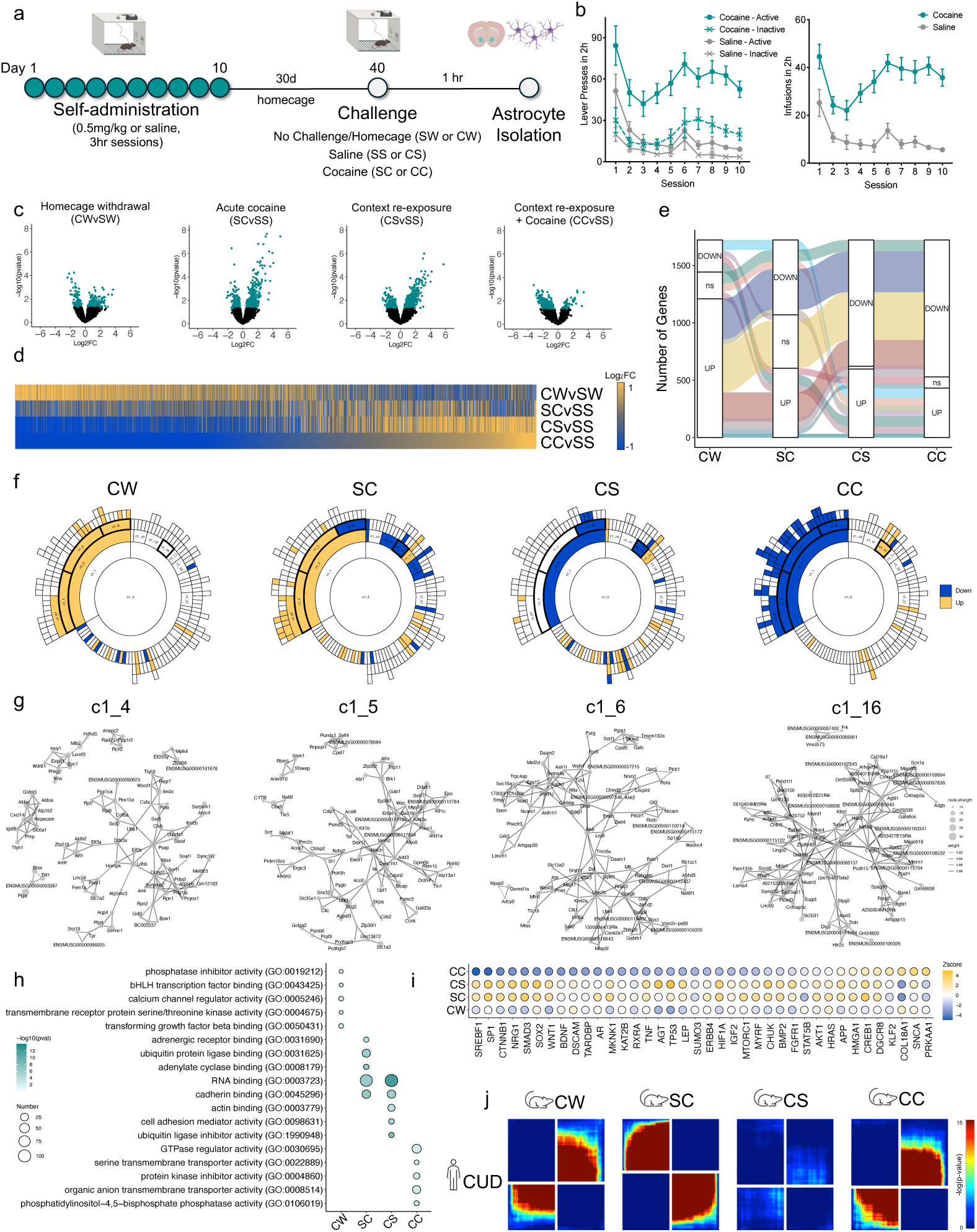
NAc astrocyte-specific RNA-sequencing after cocaine self-administration. A) Schematic illustrating experimental design and timeline for SA and astrocyte isolation for RNA-Seq. B) Acquisition of cocaine SA: mice pressed the active lever more (left; three-way ANOVA F _(1, 44)_ = 7.036, *P* < 0.05) and took more infusions than their saline counterparts (right; two-way ANOVA F _(9, 265)_ = 2.927, *P* = 0.0025). C) Volcano plots of detected genes. Significant DEGs (Log_2_FC ± 0.2 and *p* < 0.05) are indicated in teal. D) Union heatmap of Log_2_FC of significant DEGs across SA conditions, where upregulated DEGs are indicated in yellow and downregulated DEGs are indicated in blue. E) Alluvial plots demonstrate context-specific transcriptional signatures in astrocytes. F) Sunburst plots of MEGENA networks reveal similar enrichment for up- and downregulated DEGs as D and E. G) MEGENA gene networks highlight top 100 co-regulated genes where size of the bubble indicates node strength and line width indicates weight. H) Top ranked GO terms across cocaine SA conditions where size of bubble indicates number of genes and color indicates *p* value. I) Bubble plot of overlapping Upstream Regulators. Activated regulators are indicated by yellow and inhibited by blue. J) RRHO2 comparison of human CUD bulk RNA-Seq of NAc to astrocyte cocaine SA demonstrates convergent signatures between CW and CC but divergent for SC and no apparent overlap for CS.

Volcano plots demonstrate a robust transcriptional response in NAc astrocytes across all conditions, with the lowest number of differentially expressed genes (DEGs; p-value < 0.05 and Log_2_FC ± 0.2) observed in CW and the largest in CC conditions (Fig 1C and SFig1A). Both SC and CS demonstrated largely upregulated DEGs (Fig 1C and SFig1A). Interestingly, we observed that over 90% of CC DEGs were downregulated. In concordance with previous literature, we observed downregulation of *Slc1a2* and *Slc7a11*, and upregulation of *Gfap* ^23^. Union heatmap visualization revealed similar patterns of DEG expression between CC and CS, little overlap with SC, and generally opposing regulation with CW (Fig 1D). Similar DEG patterns were observed in alluvial plots (Fig 1E).

Sunburst plots of multiscale embedded gene co-expression network analysis (MEGENA) reveal modules of co-regulated genes enriched for DEGs (Fig 1F). In particular, modules c1_4 and c1_5 were enriched in upregulated DEGs in CW and SC while downregulated in CC. Module c1_6 was uniquely associated with upregulated withdrawal DEGs, while c1_16 was uniquely enriched in upregulated CC DEGs. Network plots reveal genes and their correlations with the above modules (Fig 1G). Noteworthy hub genes include *Aqp4*, *Ldhb*, and *Slc6a1* in c1_4, *Slc1a2* in c1_5, and *Ncam1*, *Nrxn1*, and *Gli3* in c1_6. Furthermore, degPattern analysis determined 9 clusters for patterns of gene abundance across cocaine SA (SFig 1E).

Venn diagrams demonstrate that despite overlapping patterns of expression (Fig 1 D-F) little overlap was observed when directly comparing DEGs across cocaine SA (SFig 1B-D). Unsurprisingly, therefore, gene ontology (Biological Process) analysis of DEGs revealed unique pathways implicated across cocaine SA conditions, largely associated with transporter activity in CC, actin and cell adhesion activity in CS, enzymatic activity in SC, and calcium and kinase activity in CW (Fig 1H). Overlapping GO terms for RNA binding and cadherin binding were observed in SC and CS conditions. To determine potential regulators specifically of identified DEGs, Upstream Regulator analysis was performed and identified several significant predicted upstream regulators including SREBF1, SMAD3, and CREB1 (Fig 1I), which were identified in a previous bulk RNA-Seq study ^24^. Finally, Rank-Rank Hypergeometric Overlap (RRHO2) analysis was performed to examine genome-wide, threshold-free astrocytic transcriptional signatures across cocaine SA conditions and human CUD ^25^. Convergence was observed between CW, CS, and CC, with divergent signatures seen for SC (SFig 1F). Importantly, RRHO2 revealed convergence of the astrocyte transcriptome with human CUD specifically for CW and CC cocaine SA and divergence for CUD and SC (Fig 1J).

### CREB is a cocaine-induced astrocyte transcriptional regulator in the NAc

Transcription factor CREB1 was identified as a predicted astrocyte upstream regulator across cocaine SA conditions, and we recently identified that CREB activation in NAc astrocytes promotes stress susceptibility ^26^. We therefore next investigated if cocaine similarly induces astrocytic CREB activation. Male mice were I.P. injected with saline or cocaine (20 mg/kg) daily for 7 d followed by either short-term (24 hr) or long-term (30 d) forced abstinence (Fig 2A). Mice were randomly assigned to receive a challenge I.P. injection of saline (SS), cocaine (SC and CC), or no injection (S and C), and 1 hr later perfused (Fig 2A). Immunohistochemistry revealed increased levels of activated CREB (pCREB, phosphorylated at serine 133) in astrocytes (Sox9+ cells) in 24C and 24CC, and a trending increase following a single cocaine injection (Fig 2B,C). After 30 d forced abstinence, we observed increased activation of astrocytic CREB in our cocaine challenge conditions (C30C), including an acute, single cocaine injection (S30C), but no change following withdrawal (30C) (Fig 2C). These data corroborate our upstream regulator analyses that CREB is indeed activated in astrocytes in response to cocaine.

**Figure 2.**
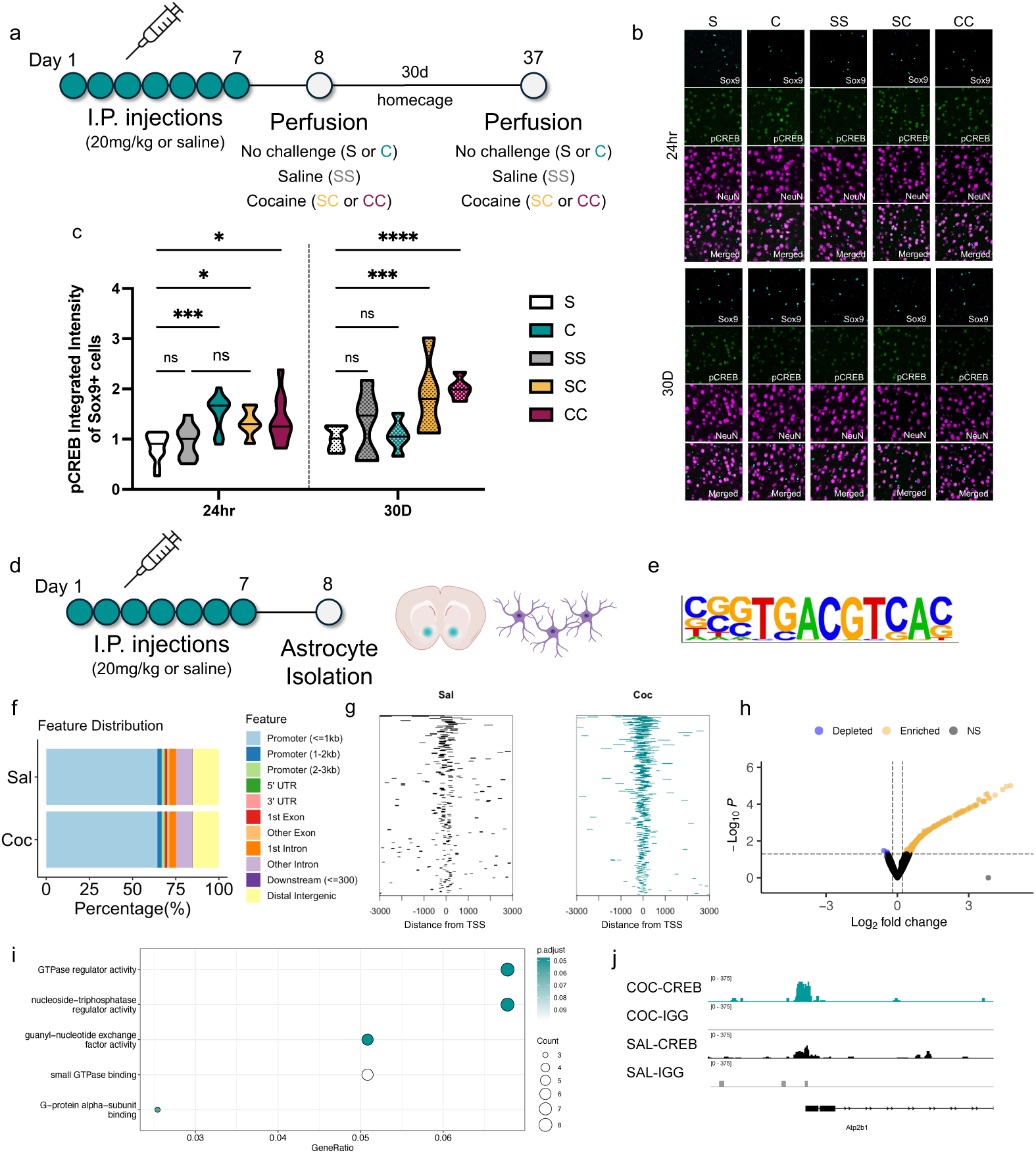
NAc astrocytic CREB is a cocaine-induced transcriptional regulator. (A) Schematic illustrating experimental design and timeline for I.P. injections. (B) Representative IHC images for pCREB (green) in astrocytes (Sox9, cyan) and neurons (NeuN, magenta) across cocaine I.P. injections (40x magnification). (C) Quantification of Sox9+ pCREB integrated intensity following 24 hr (left) or 30 d abstinence (right; two-way ANOVA interaction F_(4, 94)_ = 5.523, *P* = 0.0005). Increased astrocytic CREB activation was observed following 24 hr withdrawal from 7 d cocaine injections (C, teal, Tukey’s *p* = 0.001), with a single cocaine injection (SC, yellow, Tukey’s *p* = 0.04), and with a cocaine challenge injection (CC, magenta, p = 0.0256) compared to saline controls. Following 30D abstinence, activated astrocytic CREB was observed with a single cocaine injection (Tukey’s *p* = 0.0002) and a cocaine challenge (Tukey’s *p* < 0.0001) compared to saline controls. (D) Schematic illustrating experimental design and timeline for CUT&RUN. (E) HOMER motif analysis showing consensus CREB motif by CUT&RUN in astrocytes. (F) Genomic distribution and (G) heatmap of MACS2 called CREB peaks in astrocytes of saline- and cocaine-injected mice. (H) Volcano plot of differentially identified peaks reveals largely peaks enriched for CREB in cocaine-injected astrocytes (I) GO analysis of cocaine-induced astrocytic CREB binding sites. (J) Representative IGV tracks of CREB binding at *Atp2b1*. (IHC: n = 8-13 per condition; C&R: 2 animals pooled per biological replicate, n = 3-4 biological replicates; *p<05; ***p<0.01; ***p<0.0001)

To determine the putative gene targets of astrocytic CREB, we turned to CUT&RUN followed by DNA Sequencing (C&R-Seq)^27^. Astrocytes were acutely isolated from the NAc following 7 daily injections (I.P.) of saline or cocaine with 24 hr withdrawal (20 mg/kg; Fig 2D). HOMER motif analysis identified the CREB consensus sequence (Fig 2E). ChIPSeeker gene annotation of MACS2 called peaks revealed CREB binding predominantly at promoter regions (Fig 2F,G) in astrocytes under both saline and cocaine conditions. DiffBind analysis revealed predominantly differentially enriched CREB binding in cocaine-injected astrocytes (Fig 2H). GO analysis of enriched peaks identified biological pathways largely associated with kinase, GTPase and GPCR activity (Fig 2I). Notably, we observed increased binding of CREB at *Atp2b1*, a plasma membrane Ca^2+^ transporting ATPase, in astrocytes from cocaine-treated mice (Fig 2J).

### Astrocytic CREB in the NAc modulates cocaine reward and reinforcement

Given the well-established role of neuronal CREB in NAc in regulating addiction-related behaviors ^28,29^, we next probed if astrocytic CREB similarly modulates cocaine’s rewarding or reinforcing effects. We leveraged CREB-floxed mice in combination with viral-mediated gene transfer to bidirectionally alter CREB expression, including a previously published AAV to overexpress astrocytic CREB (CREB-OE) and an astrocyte-specific AAV driving Cre recombinase to excise CREB, both selectively in NAc astrocytes (CREB-KO; SFig 2A-D) ^26,30^. Male and female mice underwent stereotaxic surgery and then 4-5 weeks later were subjected to cocaine conditioned place preference (CPP) using a biased design (Fig 3A). Males with CREB-OE in NAc astrocytes exhibited a heightened preference for the cocaine-paired chamber compared to EGFP controls. Conversely, CREB-KO males exhibited decreased preference for cocaine (Fig 3B). This effect was only observed in males, with neither overexpression nor knockout of CREB altering CPP scores in females. To test if astrocytic CREB responds to cocaine in females, we I.P. injected females with saline or cocaine (20 mg/kg) daily for 7 days and perfused after 24 hr withdrawal (SFig 2E). We observed no difference in pCREB levels in female NAc astrocytes, suggesting that astrocytic CREB is not a cocaine-induced transcriptional regulator in females (SFig 2F,G). These results together indicate that astrocyte CREB bidirectionally influences the rewarding effects of cocaine, but in males only.

**Figure 3.**
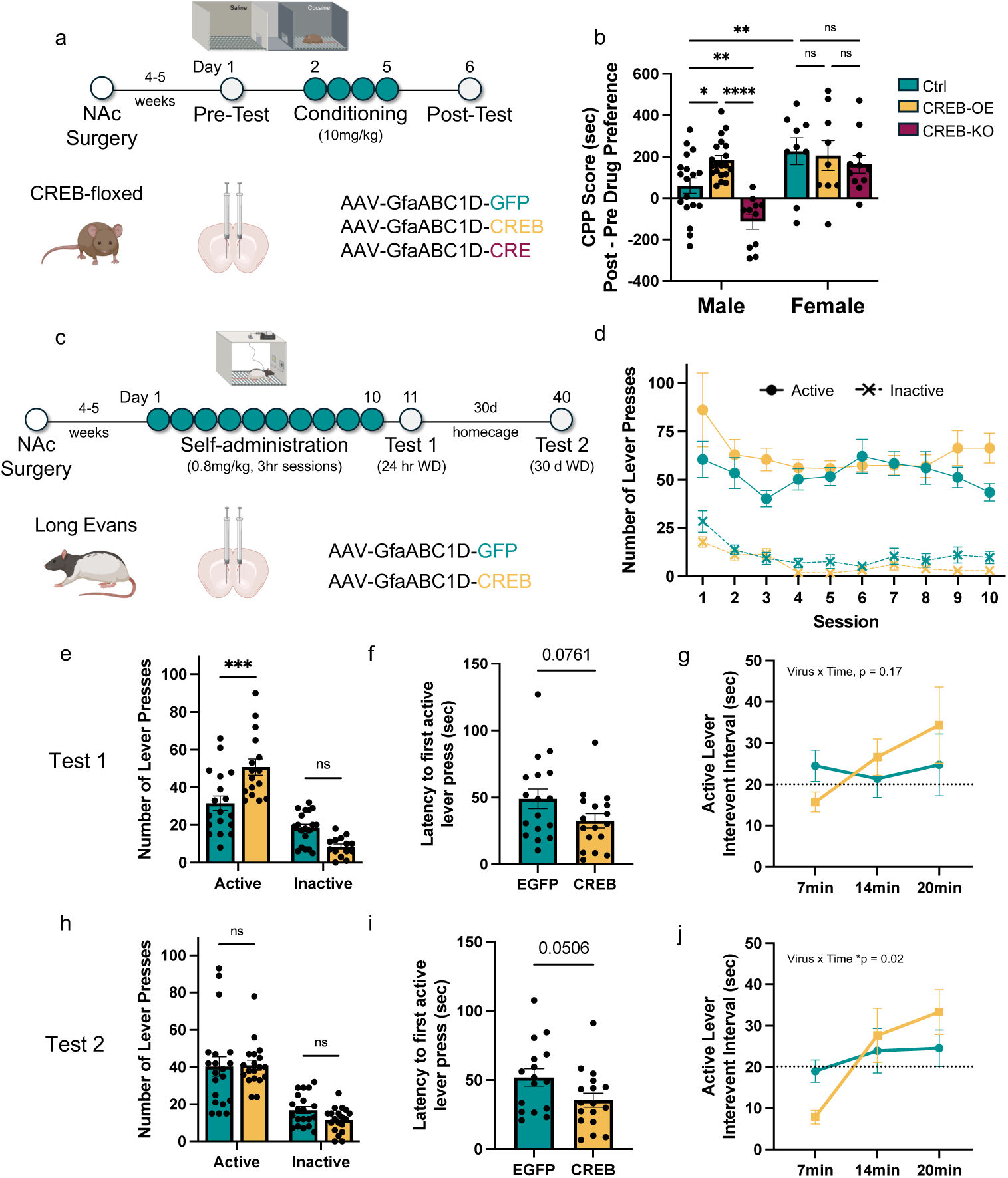
NAc astrocytic CREB regulates the rewarding and reinforcing effects of cocaine. (A) Schematic illustrating experimental design and timeline for CPP. (B) CPP score for male mice demonstrate increased preference for the cocaine-paired chamber with astrocytic CREB-OE (yellow; Tukey’s *p* = 0.0336) and a decreased preference with CREB-KO (magenta; Tukey’s *p* = 0.0091) compared to controls (Ctrl, teal) (two-way ANOVA F_(2, 72)_ = 4.3, *P* = 0.0167). No change in CPP scores was observed in females. (C) Schematic illustrating experimental design and timeline for SA. (D) Acquisition of SA demonstrates that both CREB-OE (yellow) and Ctrl (teal) rats correctly discriminated between the active and inactive levers (three-way ANOVA F_(1, 36)_ = 281.6, *P* < 0.001). Effect of CREB-OE on (E) number of lever presses (two-way ANOVA interaction F_(1, 63)_ = 20.24, *P* < 0.001), (F) latency to first lever press (t-test(32) = 1.833, *p* = 0.0761) and (G) interevent interval for active lever press (two-way ANOVA interaction F_(2, 164)_ = 1.782, *P* = 0.172) for 24 hr drug-seeking test (Test 1). Effect of CREB-OE on (H) number of lever presses (two-way ANOVA interaction F_(1, 63)_ = 0.8769, *P* = 0.3521), (I) latency to first lever press (t-test(32) = 2.034, *p* = 0.0506), and (J) interevent interval for active lever press (two-way ANOVA interaction F_(4, 232)_ = 2.95, *P* = 0.021) for 30 d abstinence drug-seeking test (Test 2). These analyses demonstrate increased drug-seeking behavior in CREB-OE rats. (n = 12-20 male and 9-11 female mice for CPP; n = 19-20 male rats for SA; Tukey’s post-hoc *p<05; ***p<0.01; ***p<0.0001).

We next tested the role of astrocytic CREB in cocaine reinforcement and seeking behavior with SA. Male Long Evans rats were stereotaxically injected with CREB-OE or EGFP-control AAV in the NAc and 4-5 weeks later jugular vein catheters inserted. Rats self-administered cocaine for 10 d on a short-access schedule (0.8 mg/kg/infusion, 3 hr sessions) followed by two 30 min drug-seeking tests wherein context cues (levers/cue lights) remain, but with no infusions (Fig 3C). Test 1 (T1) occurred 24 hr after the last SA session and Test 2 (T2) occurred after 30 d homecage abstinence. During acquisition, both EGFP and CREB-OE rats discriminated between the active and inactive levers (Fig 3D; SFig 2I). On average across all ten SA sessions, CREB-OE animals consumed more cocaine compared to EGFP controls (SFig 2J,K). During T1, CREB-OE animals exhibited increased drug-seeking behavior as measured by an increase in the number of active lever presses (Fig 3E). After 30 d forced abstinence (T2) this effect was lost, with no difference in the number of active or inactive lever presses (Fig 3H). At both T1 and T2, the total number of combined lever presses did not differ (SFig 2L,M).

We performed additional analyses to determine if 30 d abstinence decreased drug-seeking behavior in CREB-OE animals, including latency to lever press and time between active lever presses (interevent interval). During T1, CREB-OE animals demonstrated a trending decrease in latency to first active lever press (Fig 3F). Interevent intervals were binned across the session, and we observed, albeit nonsignificant, that CREB-OE animals demonstrated decreased interevent intervals at the beginning of the test with increasing intervals at later periods (Fig 3G). In contrast, EGFP controls maintained an interevent interval consistent with the timeout period (20 seconds). In T2 we again observed a trending decrease in latency to first active lever press (Fig 3I). CREB-OE animals exhibited a significant AAV x Time interaction with decreased interevent intervals at early stages of the session during T2 (Fig 3J). These data combined suggest that astrocytic CREB increases cocaine SA as well as seeking after both acute and prolonged abstinence.

### Astrocyte Ca^2+^ transients are modulated by cocaine and CREB in the NAc

Previous work implicates dynamic modulation of astrocytic calcium in response to cocaine. To date no publication has examined *in vivo* astrocyte calcium responses across repeated cocaine exposure nor has a molecular mechanism been determined ^5,7^. We therefore turned to astrocyte fiber photometry in our CREB-floxed line in combination with CPP. Male mice underwent stereotaxic surgery and fiber optic placement and then 4-5 weeks later were subjected to cocaine CPP (Fig 4A). We recorded astrocyte transients during day 1 of conditioning (acute [D1]), the last conditioning day (repeated [D4]), and during the post-test (Fig 4A,B). In response to a single cocaine injection, astrocyte Ca^2+^ transient frequency increased in control (Ctrl) astrocytes (Fig 4C). CREB-OE astrocytes exhibited a blunted effect, resulting in a nonsignificant increase in number of transients in response to cocaine (Fig 4C). CREB-KO completely abolished this effect and resulted in no change in transient number (Fig 4C). After repeated exposure, we observed no significant increase in transient frequency regardless of viral manipulation (Fig 4D). A statistically significant interaction between injection (coc vs sal) x AAV x Time suggests that in control astrocytes we observed an initial increase in Ca^2+^ transient frequency that decreases over repeated exposure which is blunted in CREB-OE and completely lost in CREB-KO astrocytes (Fig 4E). We saw no effect of cocaine or virus on transient amplitude at either timepoint (Fig 4F-H).

**Figure 4.**
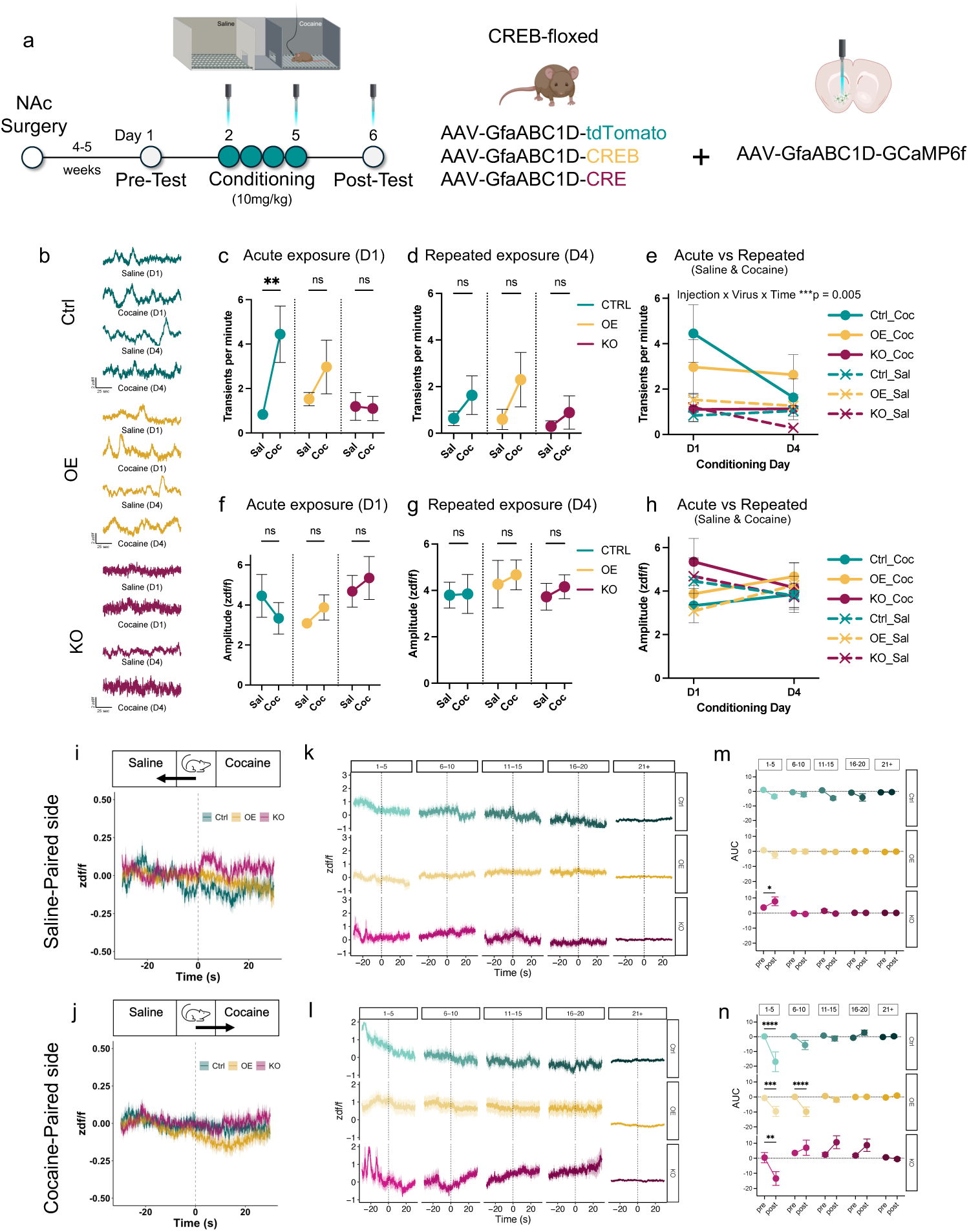
NAc astrocyte Ca^2+^ transients are modulated by cocaine and CREB. (A) Schematic illustrating experimental design and timeline for calcium fiber photometry. (B) Representative traces of astrocyte transients across viral manipulations and saline/cocaine injections. Astrocyte transient frequency in response to (C) initial (two-way ANOVA F_(1, 9)_ = 6.937, *P* = 0.0272) and (D) repeated (two-way ANOVA F_(1, 9)_ = 3.26, *P* = 0.1139) cocaine exposure separated by viral manipulation. (E) Across repeated cocaine (solid lines), astrocyte transient frequency decreases in control (Ctrl) astrocytes (teal), is blunted by CREB-OE (yellow), and not responsive to cocaine in CREB-KO (magenta; three-way ANOVA interaction F_(2, 16)_ = 7.517, *P* = 0.005). Neither cocaine nor viral manipulation affected the amplitude for (F) initial (two-way ANOVA F_(2, 10)_ = 2.68, *P* = 0.117), (G) repeated (two-way ANOVA F_(2, 10)_ = 0.0492, *P* = 0.952), or (H) across exposure (three-way ANOVA interaction F_(2, 20)_ = 0.3822, *P* = 0.6872). Representative traces averaged across the post-test as the animal enters the (I) saline or (J) cocaine-paired chambers and when binned for number of (K) saline-paired or (L) cocaine-paired entries. AUC of Ca^2+^ transients binned by chamber entry reveal that (M) CREB-KO significantly increased on initial entries to the saline-paired chamber (three-way ANOVA interaction F_(4, 280)_ = 5.403, *P* = 0.0003). (N) All astrocytes decreased Ca^2+^ AUC upon entry to the cocaine-paired chamber (Ctrl: three-way ANOVA interaction F_(4, 280)_ = 3.726, *P* = 0.0057; CREB-OE: three-way ANOVA interaction F_(4, 280)_ = 2.8, *P* = 0.0263). Only the CREB-OE astrocytes demonstrated prolonged decreased AUC in cocaine-paired chamber. (n = 5 mice per condition; Tukey’s post-hoc *p<05; ***p<0.01; ****p<0.0001).

We next examined astrocyte Ca^2+^ dynamics when the animal enters the CPP chambers during the post-test (Fig 4I,J). Averaged across the whole post-test, we observed a decrease in area under the curve (AUC) in control astrocytes when entering either saline- or cocaine-paired chambers, a decrease in CREB-OE only for cocaine-paired chambers, and CREB-KO increased Ca^2+^ after mice entered the saline-paired chamber (Fig S3G). These observations did not persist across the entire session, when Ca^2+^ dynamics were binned across the number of chamber entries (Fig 4K,L). No significant changes in AUC were found in control or CREB-OE when entering the saline-paired chamber across the binned session (Fig 4M). CREB-KO AUC increased during the first 5 entries to the saline-paired chamber followed by a return to baseline (Fig 4M). For the cocaine-paired chamber, control, CREB-OE, and CREB-KO exhibited decreased AUC upon the first 5 entries (Fig 4N). However, control astrocytes returned to baseline by 10 entries, whereas CREB-OE remained decreased through 10 entries. In contrast, CREB-KO astrocytes increased AUC after entering the cocaine-paired chamber compared to both control and CREB-OE (Fig 4N).

### Astrocytic CREB in the NAc selectively modulates D1-MSNs

Astrocyte Ca^2+^ signaling has been linked to their regulation of neuronal activity, and increased research highlights astrocytes’ ability to selectively regulate subtypes of neurons ^2,31^. Therefore, we examined if astrocytic CREB regulates the activity of major neuronal subtypes in the NAc. Male D1-tdTomato fluorescent reporter mice underwent stereotaxic surgery and then 3 weeks later whole-cell voltage-clamp recording was performed in NAc slices to measure EPSCs and IPSCs in neurons with and without fluorescence, presumably D1- and D2-MSNs, respectively (Fig 5A,B). Astrocytic CREB-OE increased the amplitude, but not frequency, of spontaneous EPSCs in D1-MSNs adjacent to the infected astrocyte processes (Fig 5C,D). These effects were not obtained in D2-MSNs (Fig 5E,F). Furthermore, neither the amplitude nor frequency of spontaneous IPSCs in D1- or D2-MSNs was affected by astrocytic CREB-OE (Fig 5G-J). This selective regulation suggests regulation of glutamatergic synaptic transmission to NAc D1-MSNs as one of key cellular substrates for astrocytic CREB signaling.

**Figure 5.**
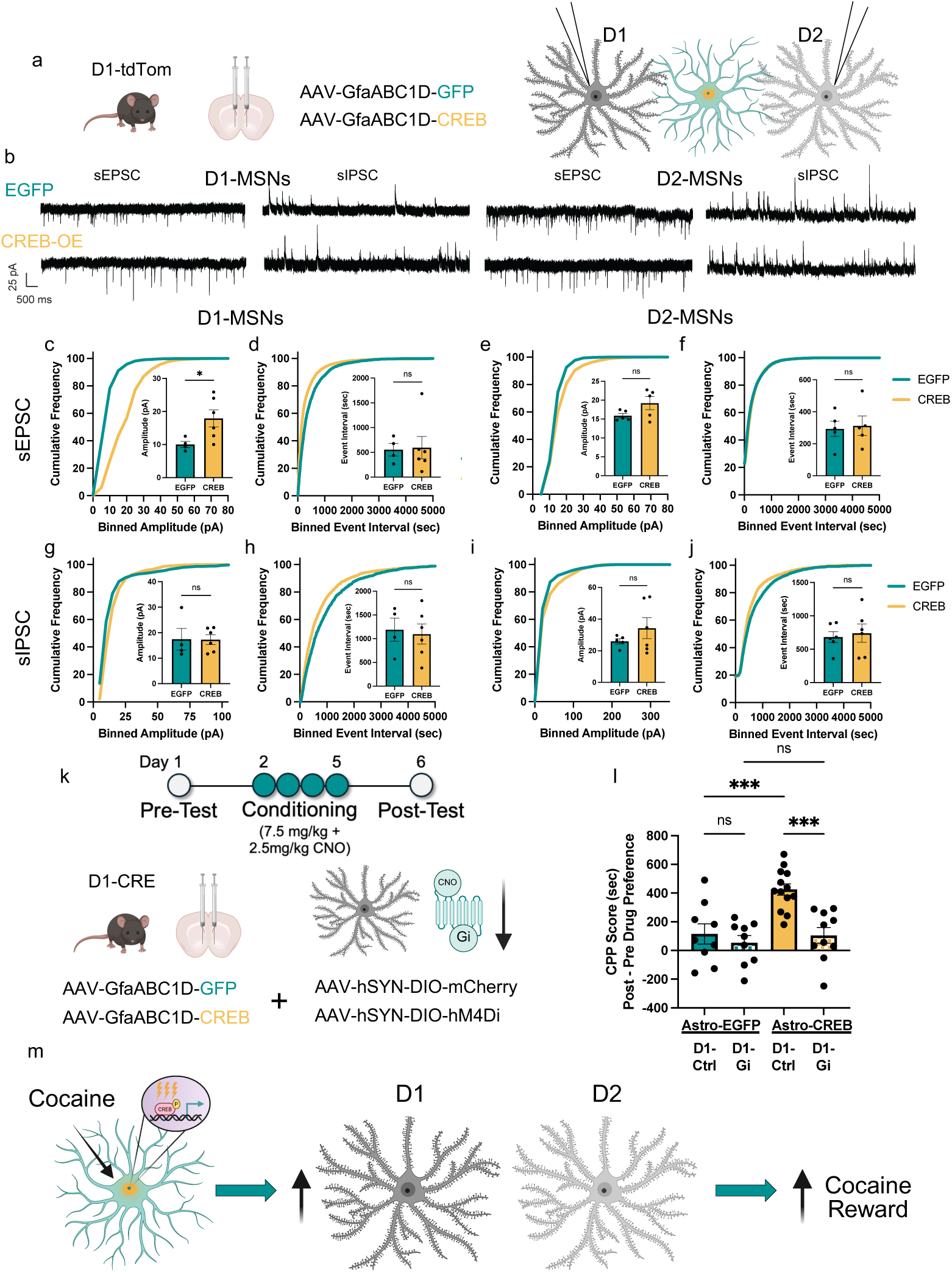
NAc astrocytic CREB selectively modulates D1-MSNs to increase cocaine reward. (A) Schematic illustrating experimental design for whole-cell patch electrophysiology experiments. (B) Representative traces of sEPSCs and sIPSCs in D1 (left) and D2-MSNs. Astrocytic CREB-OE (yellow) selectively increased excitatory synaptic inputs to D1-MSNs with (C) increased sEPSC amplitude (t(8) = 2.354, *p* = 0.046) with no influence on (D) frequency (t(8) = 0.1384, *p* = 0.8934), and no effect on sIPSC (E) amplitude (t(8) = 0.0391, *p* = 0.969) or (F) frequency (t(8) = 0.283, *p* = 0.7846) compared to control (Ctrl, teal). Astrocytic CREB-OE did not change sEPSC (G) amplitude (t(8) = 1.833, *p* = 0.104) or (H) frequency (t(8) = 0.245, *p* = 0.8123) and sIPSC (I) amplitude (t(8) = 1.115, *p* = 0.294) or (J) frequency (t(8) = 0.363, *p* = 0.724) in D2-MSNs. (K) Schematic illustrating experimental design and timeline for Gi-DREADD mediated inhibition of D1-MSNs during CPP. (L) CPP scores indicate that co-administering CNO and cocaine during conditioning (yellow/white patterned) abolishes the ability of astrocytic CREB-OE (yellow) to increase cocaine CPP. No significant effect was observed with D1-Gi-DREADD in astrocytic Ctrl animals (two-way ANOVA F_(1, 36)_ = 5.201, *P* = 0.0286) (M) Graphical abstract depicting cocaine induced activation of astrocytic CREB increases D1-MSN excitability to increase cocaine reward. (n = 4-5 animals per condition for electrophysiology; n = 10-12 animals per group for CPP; post-hoc *p<05; ***p<0.01; ****p<0.0001).

To mechanistically test if astrocytic CREB modulates addiction-related behaviors via modulation of D1-MSN activity, we coupled viral-mediated manipulations of astrocytic CREB in combination with neuronal Designer Receptors Exclusively Activated by Designer Drugs (DREADDs). Male mice underwent stereotaxic surgery and then 4-5 weeks later were subjected to CPP (Fig 5K). Importantly, CNO was administered during the cocaine conditioning sessions to silence D1-MSNs in the presence of a rewarding stimulation (cocaine), and not during the post-test. We found that Gi-DREAAD inhibition of D1-MSNs in combination with astrocytic CREB-OE resulted in loss of the cocaine-induced preference seen in controls and in astrocytic CREB-OE animals (Fig 5L). This important observation suggests that astrocytic CREB’s modulation of behavioral responses to cocaine is mediated via its selective influence on D1-MSN activity (Fig 5M).

## Discussion

Most studies of cocaine’s impact on astrocytic gene expression have relied on bulk tissue, with few directly examining genome-wide, astrocyte-specific changes^23,32,33^. Using our mouse SA model—which captures initial cocaine exposure, withdrawal/forced abstinence, context re-exposure without drug, and “relapse”/context re-exposure with drug—we identified and characterized a robust, context-specific astrocyte transcriptional response to cocaine SA. Importantly, our data converged with human CUD and replicated previously identified dysregulated NAc astrocyte genes (decreased expression of *Slc1a2*, *Slc7a11*, and *Pdgfra;* and increased expression of *Gfap* and *Pde10a*) ^23,32,33^.

We furthermore uncovered novel dysregulated astrocytic genes such as *Pla2g3* and *Etnppl*. *Pla2g3* (PLA2), which encodes phospholipase A2 group III, is one of the most significantly downregulated genes in CC and is also decreased in CUD ^34^. Astrocytic PLA2 inhibition plays a neuroprotective role in the context of glutamate toxicity; downregulation in response to cocaine re-exposure may reflect in part the glutamate hypothesis of addiction ^35^. In contrast, *Etnppl* (which encodes ethanolamine-phosphate phospholyase and is involved in lipid homeostasis and cholesterol metabolism) was upregulated in the NAc. GO analysis highlighted alterations in calcium signaling, transcription, ubiquitination, actin dynamics, GTPase and kinase activity, and serine processing—a gliotransmitter pathway linked to cocaine locomotor desensitization but not previously tied to astrocytes in CUD ^36^. MEGENA networks identified noteworthy hub genes, such as the previously identified *Slc1a2*/GLT-1 and *Aqp4*/AQP4, but also novel genes with important implications on astrocyte function such as *Ldhb*/LDHB (lactate metabolism), and *Nrxn1*/NRXN1 and *Ncam1*/NCAM1 (astrocyte-neuron synaptic interactions).

Interestingly, NAc astrocytes displayed divergent transcriptional signatures between SC and CC. Initial cocaine exposure elicited an upregulated response, whereas prolonged withdrawal and cocaine re-exposure exhibited a predominant downregulation. Previous work on bulk NAc tissue and on sorted D1- or D2-NAc MSNs revealed very different patterns, with genes upregulated in response to initial cocaine exposure exhibiting further upregulation in response to a cocaine challenge after prolonged withdrawal, an effect that dominates in D1-MSNs ^37^. In striking contrast, we observed here divergent transcriptional signatures between SC and CC in astrocytes, suggesting a fundamentally different type of reprogramming of the astrocyte transcriptome. In support of this, our predicted upstream regulators largely exhibit opposing activation status between SC and CC conditions. Notably, context exposure alone (CS) induced profound astrocyte transcriptome changes, underscoring the need for parallel neuronal data and investigation of D1/D2-MSN-dependent mechanisms.

We identified CREB as a predicted upstream regulator and determined via IHC that astrocytic CREB is activated by cocaine. We and others have implicated CREB as a transcriptional regulator in astrocytes, but direct astrocytic CREB gene targets are unknown ^26,38–41^. We therefore performed CREB C&R-Seq in acutely isolated astrocytes and determined not only increased association of CREB binding to the astrocyte genome following repeated cocaine injections, but also putative cocaine-induced gene targets. GO analysis revealed pathways such as kinase, GTPase, and GPCR activity, which correspond to GO pathways identified from our SA RNA-Seq dataset as well. Increased binding at *Atp2b1*/PMCA was particularly noteworthy given its given its role in calcium extrusion and the previously observed decrease in *ex-vivo* astrocyte Ca^2+^ transients after 14 days SA^5^.

These lines of evidence implicate astrocytic CREB as a molecular mechanism that decreases Ca^2+^ responses in astrocytes after chronic cocaine. To test this directly, we utilized fiber photometry during acute and repeated exposure, replicating that astrocyte Ca^2+^ transient frequency initially increases in response to cocaine ^5,7,42^. We extended our findings to reveal repeated exposure reduces this effect *in vivo* in the presence of cocaine. Astrocytic CREB-OE blunted cocaine-induced transient frequency—consistent with CREB-driven upregulation of *Atp2b1*/PMCA—and CREB-KO showed the opposite pattern during the post-test in the cocaine-paired chamber. These data not only implicate astrocytic Ca^2+^ signaling in bidirectional modulation of cocaine reward, aligning with prior research demonstrating Gq-DREADD attenuation of seeking and hPMCA2w/b clamping promoting cue-induced reinstatement^8,42^, but also introduce the first molecular mechanism elucidating such dynamic calcium responses in astrocytes.

Astrocytic CREB-OE rats self-administered more cocaine across acquisition and exhibited increased active lever press during the 24 hr drug-seeking test compared to EGFP controls. Although this increase in active lever pressing was not observed after 30 d, other measures of motivation and drug-seeking suggest that CREB-OE rats still exhibit increased drug-seeking behavior at this later time point. Latency to first active lever press and interevent intervals of lever presses are used as a proxy for motivation to initiate drug taking ^43–47^. The latency measure was decreased, albeit trending, at both T1 and T2 in CREB-OE rats. Reminiscent of front-loading behavior, CREB-OE rats demonstrated decreased interevent intervals at the beginning of T2 that increased as the session progressed. Together, these measures show astrocytic CREB enhances cocaine’s rewarding and reinforcing effects. Notable, neuronal CREB overexpression decreases CPP but increases SA ^28,48,49^. We reported recently that NAc astrocytic CREB increases susceptibility to social stress, similar to neuronal CREB *(22),* and optogenetic cAMP activation in hippocampal astrocytes modulates learning and memory ^50^, suggesting broader roles for astrocytic CREB in regulating complex behavior.

Female NAc astrocytes in the NAc lack typical morphological or molecular changes after prolonged withdrawal and single cell studies have revealed a sexually dimorphic response in astrocytes to a single cocaine injection ^32,51^. Consistent with this, manipulating astrocytic CREB expression in females did not affect cocaine reward. Follow up molecular approaches revealed cocaine failed to activate astrocytic CREB in females. More work is needed to disentangle the mechanisms behind such sex-specific responses.

Within the NAc, D1-MSNs are associated with positive valence and reward, while D2-MSNs generally signal aversion and extinction learning, albeit recent work suggests the contributions of these cellular populations is more complex and nuanced than this binary allocation ^52,53^. Repeated exposure to cocaine does result in bidirectional cell-type-specific changes in NAc neuronal activity that support increased cocaine reward ^54,55^. Whole cell patch electrophysiology revealed astrocytic CREB selectively enhances D1-MSN excitatory synaptic transmission, consistent with increased cocaine reward and seeking behaviors. Inhibiting D1-MSNs with Gi-DREADDs during cocaine conditioning abolished CREB-OE-induced CPP. Additionally, astrocyte Ca^2+^ signaling in the dorsal striatum decreases neuronal activity to inhibit cocaine seeking during cue-induced reinstatement, consistent with our photometry and electrophysiology findings ^42^.

Herein we demonstrate that the astrocyte transcriptome in the NAc robustly and context-specifically to cocaine SA, converging with the human CUD transcriptome. We identified CREB as a cocaine-induced astrocytic transcriptional regulator and, for the first time, show that astrocytic transcriptional regulation controls cocaine reward and reinforcement. Subsequent experiments identify potential molecular mechanisms of astrocytic CREB’s influence through modulating astrocytic Ca^2+^ signaling and selectively increasing D1-MSN activity.

## Acknowledgements

The authors would like to thank Katherine Beach, Catherine McManus, Kyra Schmidt, Nathalia Pulido, and Ezekiell Mouzon for animal husbandry; and Joseph Landry and James Callens (Yasmin Hurd lab) for technical support.

## Authors contributions

Conceptualization: L.M.H. and E.J.N. Methodology: L.M.H., C.B.J., and E.J.N. Investigation: L.M.H, A.M-T., R.F., C.J.B., F.J.M-R., T.M., T.M.G., S-Y.Y. E.M.P., M.R., C.A., Y.YY, V.K., S.Y, E.K., A.L., G.R. Visualization: L.M.H. Formal analysis: L.M.H., M.E., T.M.G, C.J.B. Resources: E.J.N. Supervision: Y.D., L.S., and E.J.N. Writing: L.M.H. and E.J.N.

---

**Figure S1.**
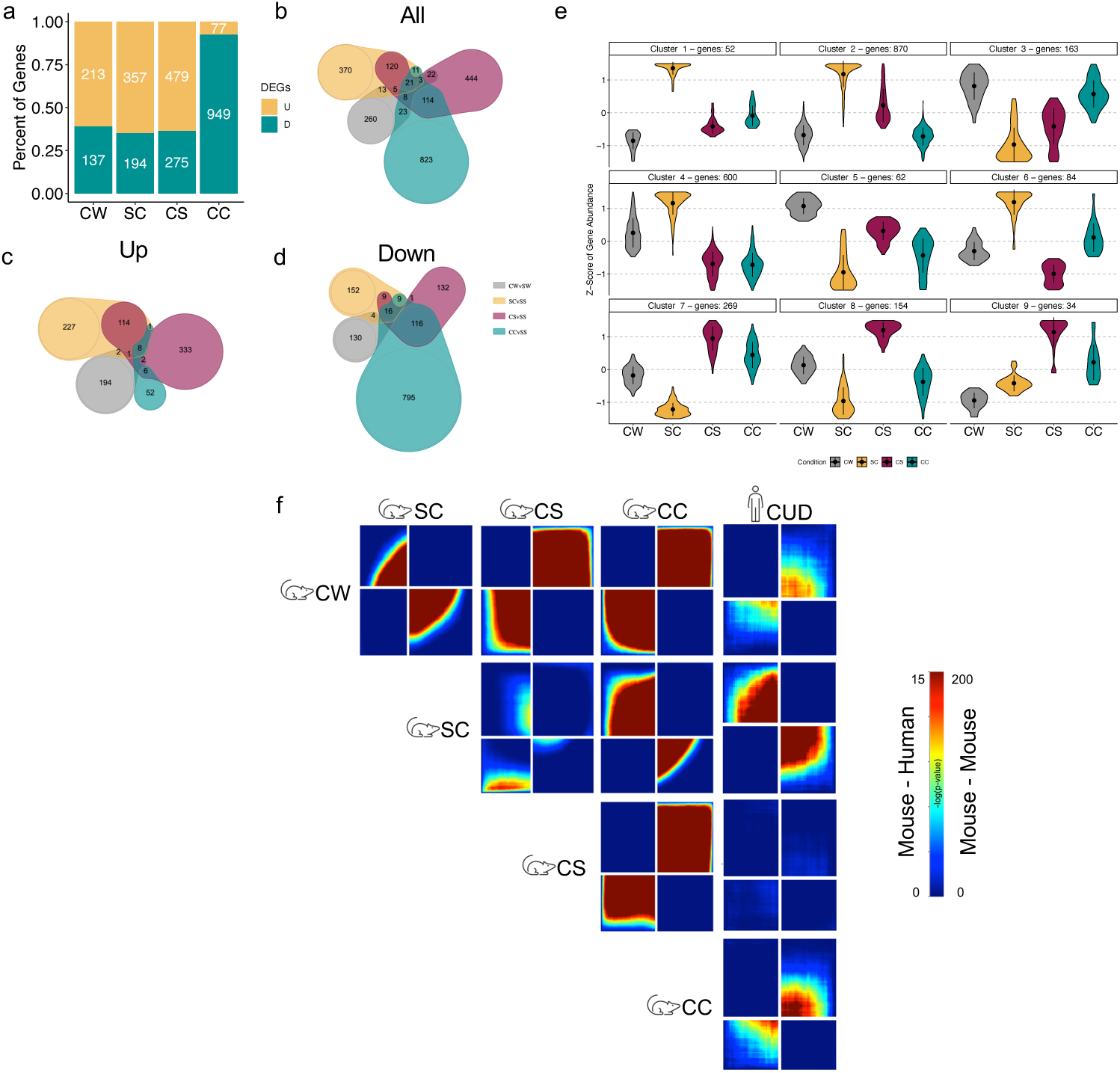
A) Percent and number of up- (yellow) and downregulated (teal) DEGs in NAc astrocytes after cocaine SA. Venn diagrams for B) all, C) up-, and D) downregulated DEGs between withdrawal (CW), acute cocaine (SC), context re-exposure (CS), and “relapse”/context re-exposure and drug re-exposure (CC), all compared to SS controls. E) ClusterProfiler identified clusters of gene abundance in astrocyte RNA-Seq after cocaine SA. F) RRHO2 comparing transcriptional signatures across mouse astrocyte RNA-Seq and human postmortem CUD bulk RNA-Seq.

**Figure S2.**
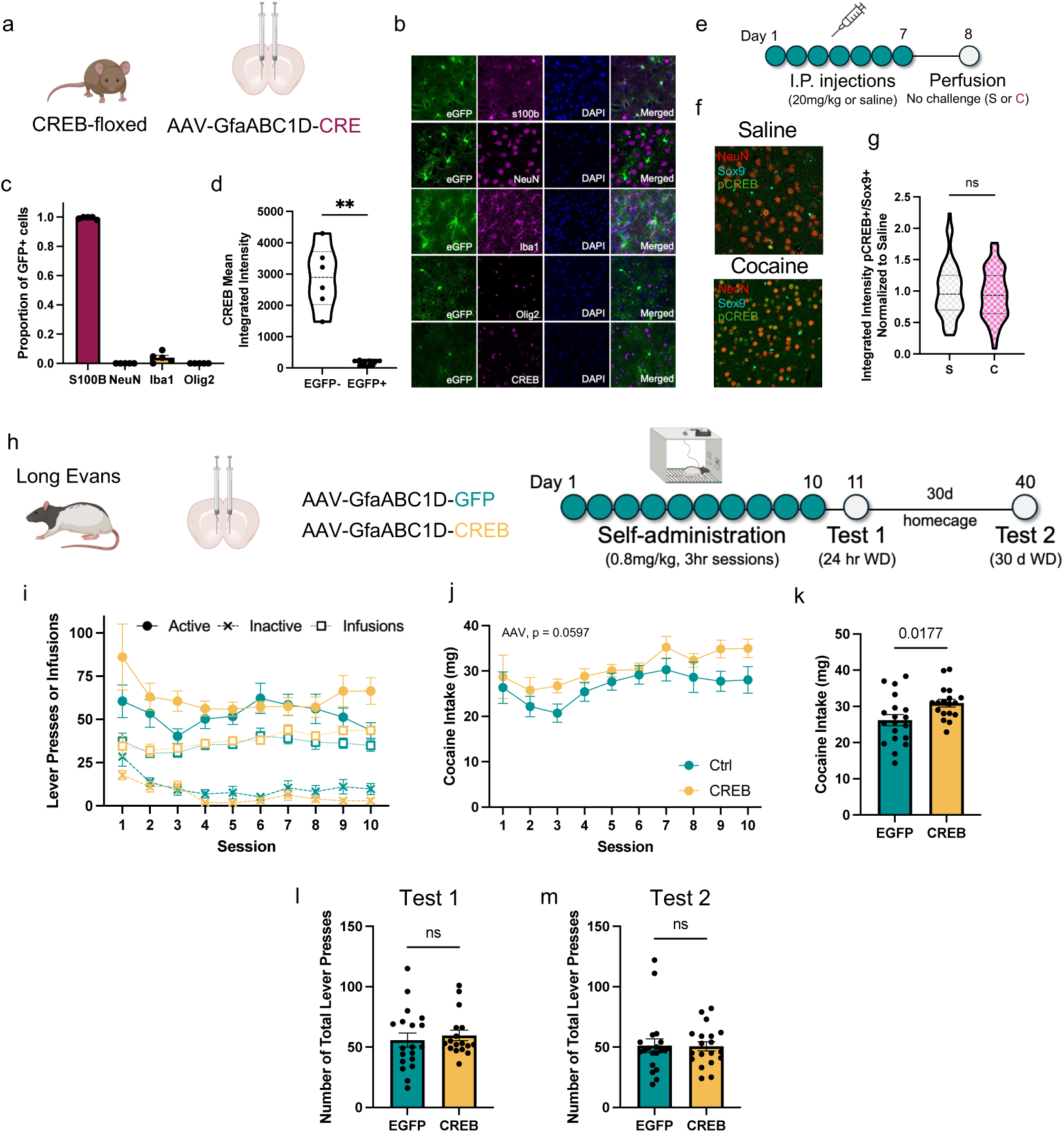
(A) Cartoon illustration of experimental design for AAV2/5-Gfap(0.7)-iCre validation. (B) Representative images of AAV-eGFP and astrocytic S100B, neuronal NEUN, microglial IBA1, oligodendrocyte OLIG2, and total CREB (all magenta) and DAPI (blue). (C) Quantification of GFP+ cells that overlap with above cellular markers demonstrates astrocyte-specific AAV expression (n = 5 animals). (D) Quantification of total CREB integrated intensity demonstrates successful excision of astrocytic CREB with AAV2/5-Gfap(0.7)-iCre in CREB-floxed mice (Welsh’s t-test: t(5.037) = 6.58, *p* = 0.0012; n = 5 animals). (E) Schematic illustrating experimental design and timeline for female I.P. injections. (F) Representative image of SOX9+ (cyan), NEUN (red), and pCREB (green) (40x magnification) for NAc of female saline- and cocaine-injected mice. (G) No change was found in activated CREB (pCREB) in female astrocytes following repeated cocaine injections (t(5) = 0.393, *p* = 0.711; n = 4-5 females per condition). H) Schematic illustrating experimental design and timeline for rat cocaine SA. I) Active (solid line/circles), inactive (dashed line/x), and infusions (dotted line/squares) for control (Ctrl, teal) and CREB-OE (yellow) rats across cocaine SA acquisition. J) Cocaine intake is trending increased in CREB-OE (yellow) rats compared to Ctrl (teal) across cocaine SA acquisition sessions (Mixed-effects F(_1, 35)_ = 3.786, *P* = 0.0597). K) Increased cocaine intake is observed when averaged per animal across all SA sessions (t(35) = 2.488, *p* = 0.017). Total number (active + inactive) lever presses for L) Test 1 (t(35) = 0.5145, *p* = 0.6102) and M) Test 2 (t(35) = 0.0.0832, *p* = 0.934).

**Figure S3.**
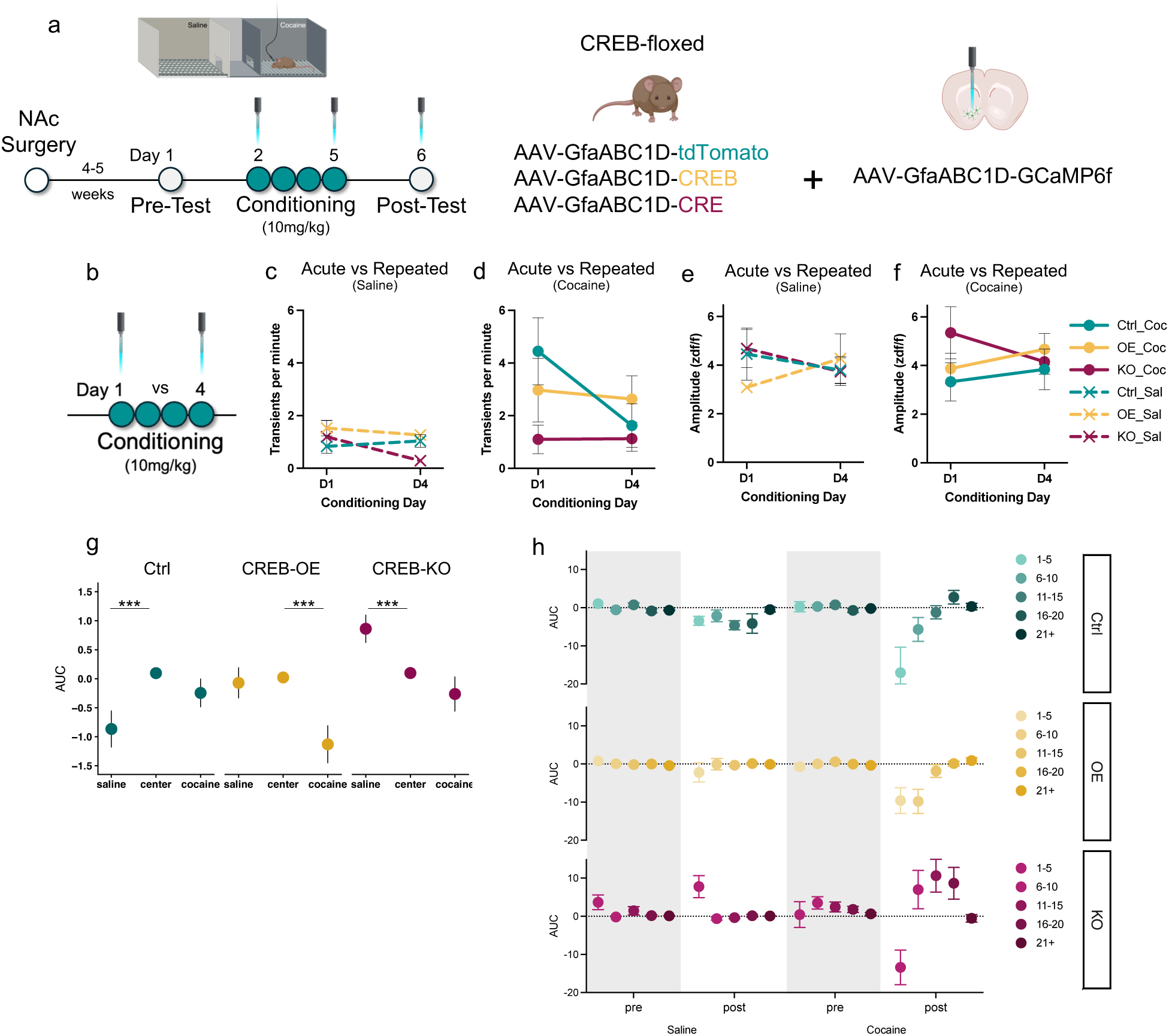
A) Schematic illustrating experimental design and timeline for fiber photometry and CPP. B) CREB viral manipulation in NAc and conditioning session comparisons of Ca^2+^ transients across days D1 and D4 conditioning sessions, including frequency for C) saline and D) cocaine and amplitude for E) saline and F) cocaine (three-way ANOVA interaction F_(2, 20)_ = 0.3822, *P* = 0.6872). G) AUC quantification of control (Ctrl, teal), CREB-OE (yellow), and CREB-KO (magenta) across the entire post-test session when entering saline-paired (left) and cocaine-paired (right) from center (middle) (three-way ANOVA F_(4, 215)_ = 9.075, *P* < 0.001; Tukey post-hoc *p<05; ***p<0.01; ****p<0.0001). H) Calcium AUC quantification of Ctrl (teal), CREB-OE (yellow), and CREB-KO (magenta) transients binned by entry number (increasing color intensity) demonstrates opposite direction of astrocytic Ca^2+^ response between CREB-KO and Ctrl and CREB-OE when entering the cocaine-paired chamber.

